# Drive, suppression, and escape from suppression of a selfish chromosome

**DOI:** 10.1101/2025.08.07.669186

**Authors:** Jackson T. Ridges, Jackson Bladen, Robert L. Unckless, Nitin Phadnis

## Abstract

Meiotic drivers are selfish chromosomes that are predicted to spark a rapid intragenomic arms-race with their suppressors. However, the long-term persistence of unsuppressed selfish chromosomes in natural populations violates these theoretical expectations. The *Drosophila pseudoobscura Sex-Ratio* (*SR*) chromosome exemplifies this problem, sometimes referred to as the “ancient gene drive paradox”. Here, we reconstruct the evolutionary history of this *SR* chromosome and show that its genetic architecture and complexity has been shaped by a history of drive, suppression, and escape from suppression. Our results indicate that the current lack of resistance to the *SR* chromosome represents a transient condition awaiting the emergence of new suppressors.

**SIGNIFICANCE:** Intragenomic arms races triggered by selfish chromosomes are expected to drive rapid evolution through cycles of drive, suppression, and escape from suppression. Despite these expectations, some selfish chromosomes such as the *D. pseudoobscura Sex-Ratio* (*SR*) chromosome exist unsuppressed for extended periods of time. Despite the absence of suppression, we uncover currently segregating suppressors against the ancestral version and evidence of recent dynamic evolution on the *SR* chromosome. Today’s *SR* chromosome thus represents a transient state in a surprisingly slow arms race. Taken together, the lack of suppressors against this selfish chromosome can be explained by mutational limits for the emergence of suppressors.

## INTRODUCTION

Meiotic drivers are selfish chromosomes that selectively prevent the development of gametes bearing their competing homolog (1–3). As such, meiotic drivers – especially those on sex chromosomes – are predicted to spark a rapid intragenomic arms-races (2). They spread through populations despite inducing fitness costs to individuals that carry them (4). Resistant target chromosomes and suppressors in other parts of the genome are expected to evolve rapidly in response to the strong selective pressure imposed by drivers (5, 6). After the driver is suppressed, new variants of the driver that escape suppression are expected to emerge, and so on, resulting in an evolutionary arms race between drivers and the rest of the genome. In violation of these strong theoretical predictions, some selfish chromosomes persist at high frequencies over long evolutionary timescales with no emergence of resistance to drive (7–9). This observed inconsistency with the framework of evolutionary arms races has been referred to as the ‘ancient gene drive paradox’ (7).

The best studied example of the ancient gene drive paradox is the *Sex-Ratio (SR)* chromosome of *D. pseudoobscura* (10). This *SR* chromosome selectively destroys *Y*-bearing sperm during spermiogenesis, causing *SR* males to yield almost entirely female offspring. Even when rare male offspring are produced, they are sterile products of non-disjunction events (*XO* males) rather than *Y*-chromosomes escaping the *SR* chromosome (11). Since its discovery in 1936, repeated genetic surveys of natural populations have shown that *SR* is consistently found at species-wide frequency of 15% (up to 30% in the American Southwest) (10, 12–17). This maintenance of the *SR* chromosome at a stable frequency indirectly suggests that no cost-free resistant *Y*-chromosomes or autosomal suppressors against *SR* have evolved during this time.

Additionally, a few direct surveys of natural populations carried out during this time have failed to detect resistant *Y* chromosomes or autosomal suppressors against *SR* (7). While factors such as female remating frequency and fitness costs to *SR*-carrying females likely explain why the *SR* chromosome has not fixed in populations and caused extinction (18, 19), the absence of genetic resistance or suppression to drive despite strong evolutionary pressure over decades remains problematic for the evolutionary arms race framework.

## RESULTS

### No suppressors of *SR* in a natural population of *D. pseudoobscura*

One possible resolution to the ancient gene drive paradox is that drive-resistant *Y*-chromosomes and autosomal suppressors exist at low frequency but went undetected in past surveys or may have emerged very recently. To explore this possibility, we conducted a systematic search for genetic suppressors against *SR* from wild-caught flies collected near Zion National Park, Utah, where *SR* is found at high frequencies. We caught ∼300 wild *D. pseudoobscura* males and crossed each male individually to laboratory females heterozygous for *SR* and *ST* (the standard non-driving X chromosome). This cross tests for drive-resistant *Y*-chromosomes or dominant autosomal suppressors of drive, as it produces sons carrying either the *ST* or *SR* chromosome and the *Y*-chromosome and a set of autosomes from the wild-caught father. Males carrying the *SR* chromosome produced from these 300 genetic backgrounds produced nearly all female offspring and internally controlled males carrying the *ST* chromosome produce typical progeny sex-ratio (Figure 1A, 1B). We also tested the strength of *SR* against an additional panel of eight strains recently collected in and near Southern Utah (gift from Steve Schaeffer). Again, males carrying the *SR* chromosome produced almost entirely female offspring, regardless of the genetic background (Figure 1C). Altogether, we find no evidence of dominant autosomal suppressors against *SR* or drive-resistant *Y*-chromosomes, even in populations where *SR* is most prevalent.

**Figure 1:**
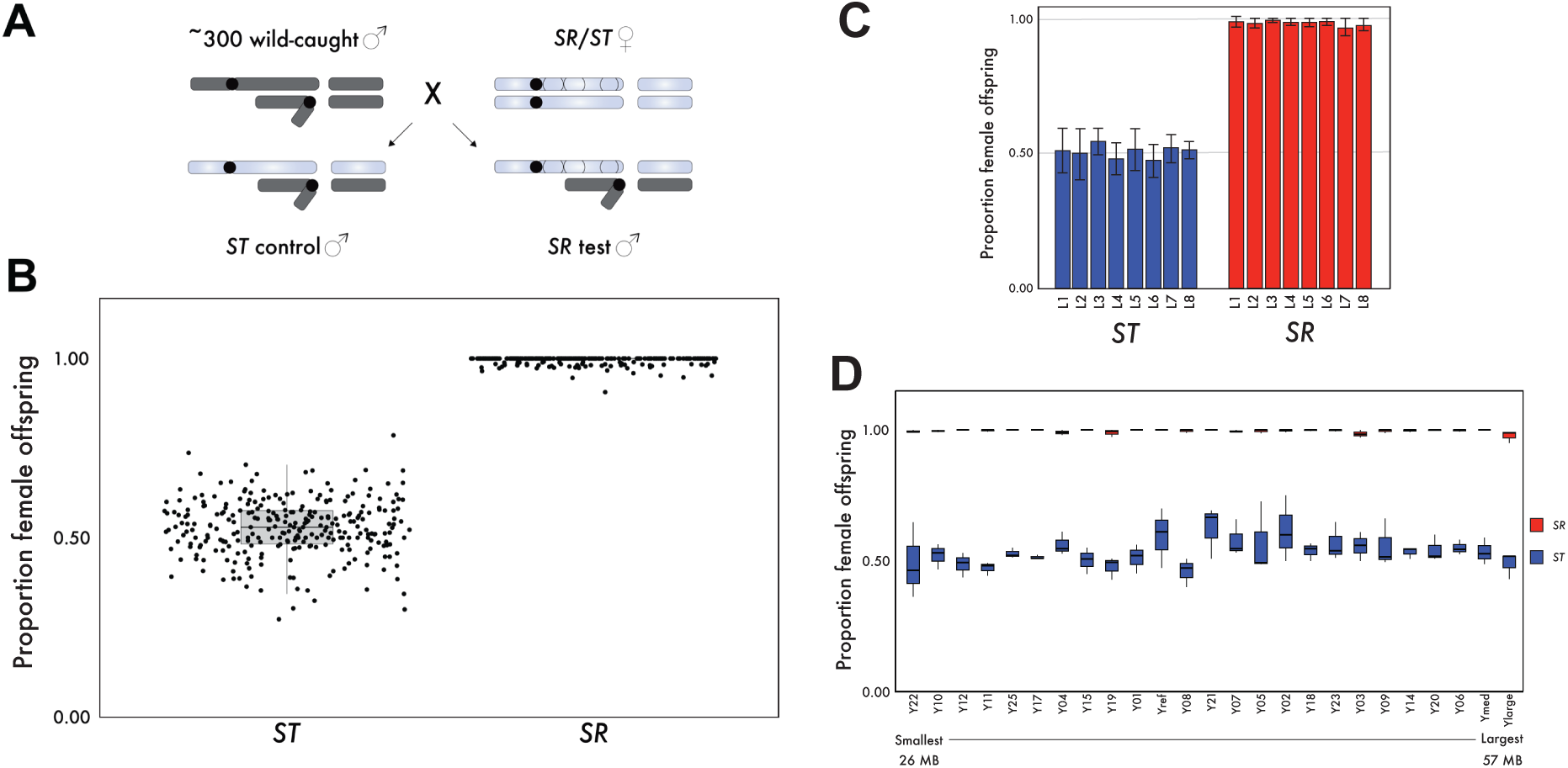
There is no evidence of suppression or resistance to the *D. pseudoobscura SR* chromosome. **(A)** Crossing scheme to test for *Y*-linked suppressors and dominant autosomal suppressors of the *D.pseudoobscura SR* chromosome. Wild-caught males are crossed to *SR/ST* females, where the standard *X*-chromosome is marked with *sepia (se)*, and *short (sh)*. This produces control *se,sh* males without the *SR* chromosome, and test *SR* males. **(B)** Progeny sex-ratio of test and control males for ∼300 wild caught *Y*-chromosomes and autosomes. No suppressors of *SR* were detected. **(C)** Females with *ST* and *SR* chromosomes were crossed to males from eight independent isogenic strains originally collected in or around southern Utah. *ST* males produce normal sex-ratios of progeny. The *SR* chromosome drives fully in all autosomal backgrounds. **(D)** *SR*/*ST* heterozygous females were crossed to a panel of males that are genetically identical except for their *Y*-chromosomes, which differ substantially in sequence, size and structure. This generated *SR* test and *ST* control males for each *Y*-chromosome, and sex-ratio of their progeny was quantified. *SR* males drive fully against all *Y*-chromosomes and *ST* males produce even sex-ratio progeny.

### Absence of *SR*-resistant *Y*-chromosomes

If *D. pseudoobscura* lacks sufficient genetic variation, specifically on the targeted *Y*-chromosome, – or if we failed to sample enough *Y*-chromosome variation from >300 wild-caught flies in Utah – this may explain our failure to detect any resistance to *SR*. Molecular evolution studies, however, have shown that *D. pseudoobscura* has a large effective population size, high levels of gene flow, and harbors substantial natural genetic variation (20–22). Cytological studies also reveal a particularly high level of variation among *Y*-chromosomes in size and structure (23–25). To detect drive resistant *Y*-chromosomes, we tested the strength of *SR* drive against a panel that captures variation on the *Y*-chromosome collected from across the species range in an otherwise isogenic background. This panel of 22 *Y*-chromosomes range in size from 26.1 MB to 56.7 MB, with a median size of 31.04 MB (gift from Doris Bachtrog). We crossed males from each *Y*-chromosome strain to females heterozygous for *SR* and *ST.* This cross produced F1 sons that carry the *ST* or *SR* chromosome and *Y*-chromosomes from the diversity panel. For every *Y*-chromosome strain, we crossed both F1 *ST* and *SR* males to *D. pseudoobscura* females and scored the sex of their offspring. The *SR* chromosome drives at nearly 100% efficiency against all *Y*-chromosomes (Figure 1D). Even though the *Y*-chromosomes assayed here vary greatly in size and sequence, we saw no evidence of drive resistant *Y*-chromosomes. These results support our assertion that genetic suppressors against *Sex-Ratio* drive are absent in *D. pseudoobscura*.

### The ancestral form of *SR* is suppressible

Another possible explanation for the lack of evidence of an arms race between the *SR* chromosome and its suppressors is that there may be an undiscovered biological quirk of *D. pseudoobscura* that limits it from evolving suppression to drive (7). We, therefore, asked whether there is any evidence for past suppression of drive to a reconstructed analog of the ancestral version of the *SR* chromosome. The *D. pseudoobscura SR* chromosome carries three non-overlapping inversions on *XR* – the *Basal* (*B*), *Medial* (*M*), and *Terminal* (*T*) inversions – that are found in complete linkage with each other in natural populations (Figure 2A). The *Basal* and *Medial* inversions of *SR* originated nearly simultaneously around one million years ago and cannot be recombinationally separated from each other. The *Terminal* inversion, however, is younger and, in rare cases, can be recombinationally separated from the *Basal*-*Medial* (*BM*) inversions (26). We have previously shown that the *BM* inversions alone are capable of sex-ratio drive, although to a weaker extent compared to the full *SR* chromosome (26). The age of the *BM* inversions along with their ability to drive on their own suggest that these two inversions represent the ancestral state of the *SR* chromosome. It is unclear whether the ancestral *BM* driver is generally weaker, or if it was previously tested in a background where it is partially suppressed.

**Figure 2:**
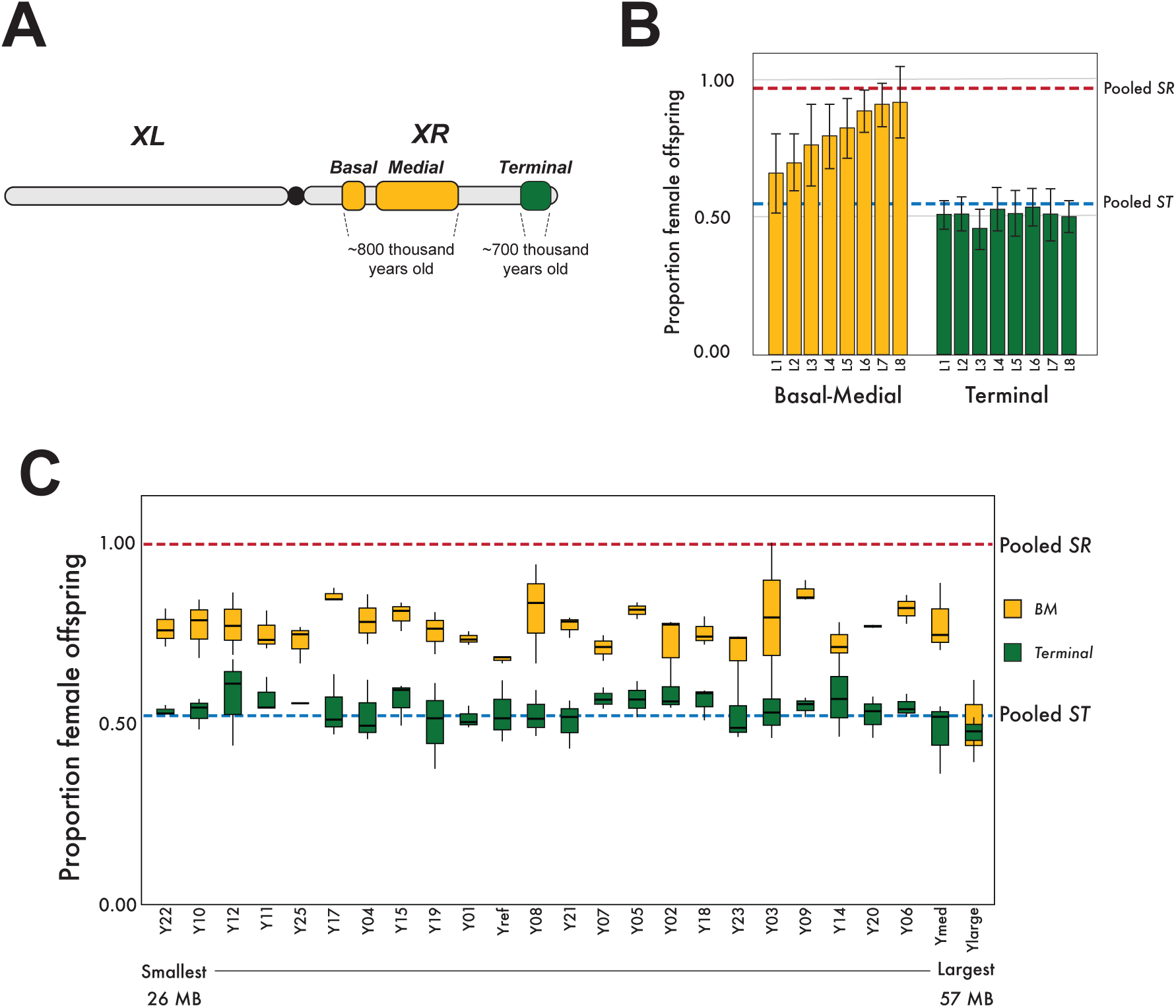
*BM* drives on its own, but is suppressed in some backgrounds and *Terminal* does not drive on its own, but enhances *BM* drive. **(A)** Schematic showing the *SR* chromosome with its three inversions on *XR*. *Basal* and *Medial* are inseparable by recombination and have been dated to ∼800,000 years old. *Terminal* is separable from *BM* through rare recombination events and has been dated to ∼700,000 years old (26). **(B)** Females carrying *Terminal* or *BM* were crossed to males from eight independent isogenic strains originally collected in or around Southern Utah. *Terminal* males produce normal sex-ratios of progeny. *BM* displays variable drive dependent on genetic background. **(C)** Males were generated that contain either the *BM* or *T* inversions over each *Y*-chromosome from the *Y*-chromosome panel, and the sex-ratio of their progeny was quantified. The *BM* inversions have variable drive depending on the *Y*-chromosome it is paired with, and *Terminal* never drives on its own regardless of *Y*-chromosome.

To test whether the *SR* chromosome has been suppressed in its evolutionary past, we measured the strength of *BM* drive against our panel of eight recently caught strains and, surprisingly, we observed substantial variation for the strength of drive (Figure 2B). We also tested the strength of *BM* drive against our *Y*-chromosome diversity panel. We observed substantial variation in the strength of *BM* drive, with lines exhibiting nearly complete drive (>95%) to near Mendelian sex-ratios (∼55%) (Figure 2C). These results show that the *BM* ancestral analog of *SR* is still capable of full drive in some genomic contexts, but drive-resistant *Y*-chromosomes and/or autosomal suppressors against the ancestral *BM* chromosome exist and still segregate in natural populations. We conclude that the *D. pseudoobscura SR* chromosome has indeed been suppressed in its past, consistent with the prediction of an evolutionary arms race.

### The *Terminal* inversion neutralizes all suppressors of the ancestral *SR* chromosome

The *Terminal* inversion is younger than the *BM* inversions and does not appear to cause sex-ratio drive alone (26). The contribution of the *Terminal* inversion to *SR* drive thus remains unclear. Based on our past findings, it is possible that the *Terminal* inversion can drive in some backgrounds, but it was previously tested in a background where it was suppressed. If *Terminal* can independently drive in some backgrounds, it may be an additive enhancer of an otherwise inefficient drive mechanism in the ancestral *SR* chromosome. Under this scenario, the *Terminal* inversion would act as an enhancer of drive in a fashion similar to *Enhancer of SD (E(SD))* in *D. melanogaster,* which can drive independently of *Segregation Distorter (SD)*, and enhances drive when *E(SD)* and *SD* are found together (27). Alternatively, changes in the *Terminal* inversion may allow the *SR* chromosome to override drive-resistant *Y*-chromosomes or counter autosomal suppressors, though *Terminal* may not be capable of driving on its own.

To distinguish between these possibilities, we crossed the *Terminal* chromosome to our *Y*-chromosome diversity and wild-caught panels (Figure 2B, 2C). We observed that the *Terminal* inversion is incapable of sex-ratio drive regardless of genetic background, including against those that are permissive to the ancestral *BM* driver. Our results show that the *Terminal* inversion does not drive on its own but interacts with the *BM* inversions through complex epistasis and converts all *BM* drive to near 100% efficiency across all genetic backgrounds. In *D. melanogaster SD*, multiple loci capable of drive on their own are locked on the same haplotype to produce strong drive. In contrast, in *D. pseudoobscura*, *Terminal* does not drive on its own yet enhances *BM* drive. The addition of *Terminal* thus enables the *SR* chromosome to drive fully, even in the presence of suppressors.

### Selective sweep at the *Terminal* inversion of the *SR* chromosome

Our results suggest that the lack of resistance against *SR* may represent a transient state where the extant version of the *D. pseudoobscura SR* chromosome currently holds the upper hand. This scenario of *drive* – *suppression* – *escape from suppression* predicts dynamic evolution of the *SR* chromosome shaped by recurrent selective sweeps on the *SR* chromosome as a part of an ongoing evolutionary arms race. To test this idea, we measured patterns of genetic diversity across *D. pseudoobscura X*-chromosomes by individually sequencing 14 *ST* and 14 *SR* wild-caught males near Zion National Park. We used PCANGSD (28) to create principal component analyses of the X chromosomes of these 28 flies (Figure 3A). Consistent with previous observations, the PCAs revealed little clustering on *XL* where recombination happens freely between *ST* and *SR* chromosomes. On *XR*, the first principal component distinguishes *ST* and *SR* chromosomes. Unexpectedly, in the *Basal* and *Medial* inversions, the *SR X* chromosomes form two discrete clusters, which suggest that there may be multiple distinct haplotypes of *BM* in natural populations. On *Terminal* however, all *SR* chromosomes cluster together tightly. One possible interpretation of this analysis is that there are multiple haplotypes of the SR chromosome at *BM*, but that each of these haplotypes carry the same *Terminal* inversion.

**Figure 3:**
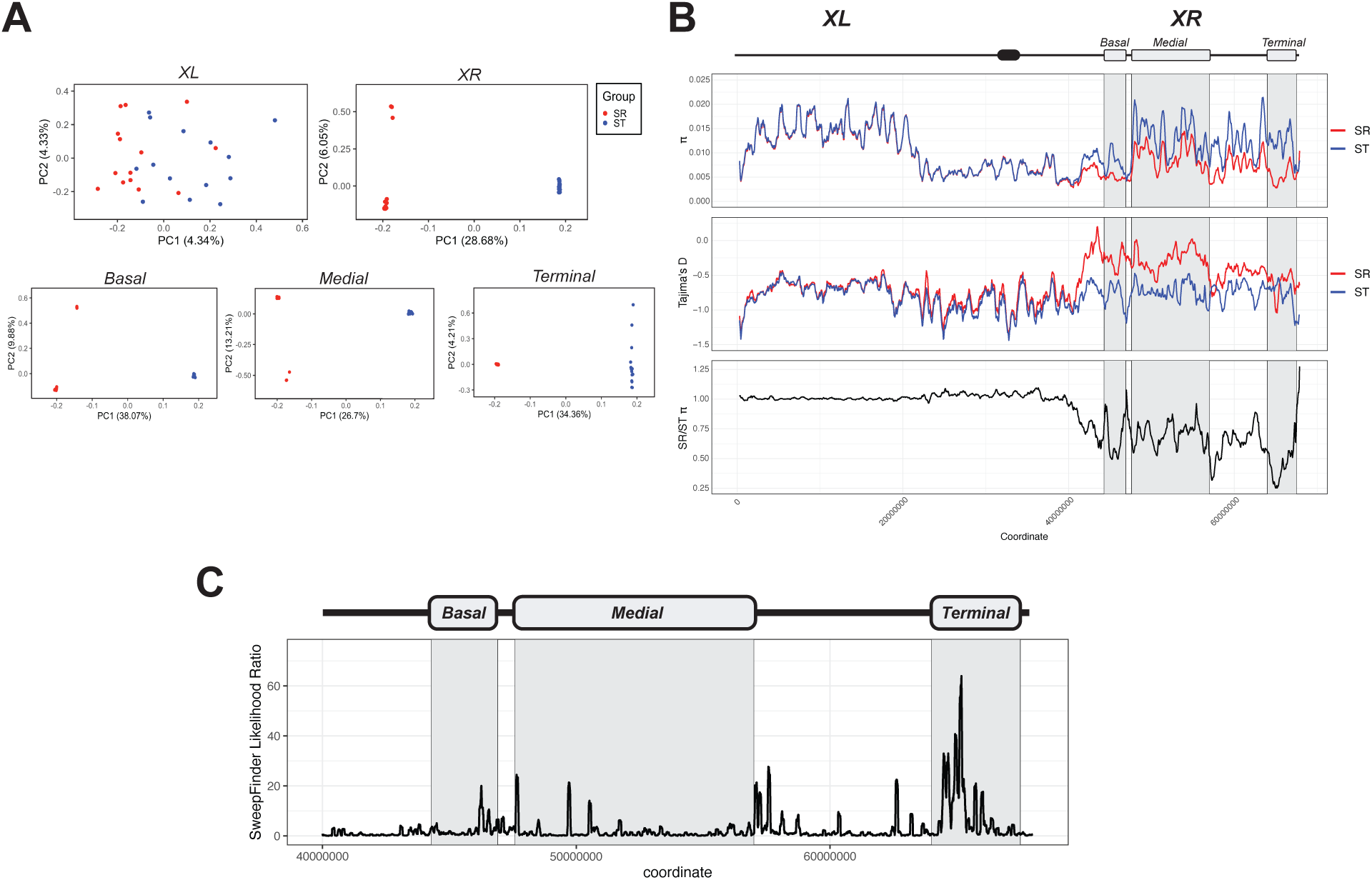
Analysis of genetic diversity across *ST* and *SR X*-chromosome reveals a recent selective sweep on *Terminal*. **(A)** vPrincipal Component Analyses of 28 wild-caught males for *XL*, *XR* and each of the three inversions on the *SR* chromosome. Individuals are colored by whether they have the *ST* or *SR X*-chromosome. At *XL*, there is no clustering of *ST* and *SR* chromosomes, consistent with previous observations that recombination suppression between *SR-ST X*-chromosomes is limited to *XR* (26). At *XR*, *SR* chromosomes cluster into two groups, indicating that there may be two distinct haplotypes of the *SR* chromosome among the 14 sequenced males. Specifically, in the *Basal* and *Medial* inversions, there again appears to be two distinct haplotypes, but the *Terminal* inversion appears to be nearly identical among sequenced SR males. **(B)** π, Tajima’s D, and *SR*/*ST* π ratio across the *X* chromosome for 28 individually sequenced *ST* and *SR* males. On *XL*, all metrics are nearly identical between *ST* and *SR* flies. On *XR*, π is generally lower on *SR* compared to *ST* but mirrors the patterns of high and low diversity, apart from at *Terminal*, where the *SR* chromosome has a large dip that the *ST* terminal does not, possibly reflecting a recent selective sweep on *SR*. **(C)** Sweepfinder2 results on *XR* for *SR* males. Sweepfinder2 predicts a large selective sweep on the *Terminal* inversion, consistent with our other analyses that indicate a depletion of genetic diversity on the *Terminal* inversion of the *SR* chromosome.

We next used ANGSD (29) to estimate π and Tajima’s D along the *X*-chromosome (Figure 3B). The *ST* and *SR* flies have nearly identical patterns of genetic diversity on the left arm of the *X*-chromosome. On *XR*, where the inversions of the *SR* chromosome reside and recombination is reduced, levels of π are generally lower on the *SR* chromosome compared to *ST* but broadly track each other across most of the chromosome arm, indicating shared population genetic forces. However, levels of *pi* are dramatically reduced at *Terminal* of *SR*, in a departure from that of the *ST* chromosome (Figure 3C). The levels of π at the *Terminal* inversion of the *SR* chromosome are among the lowest found anywhere in the genome, consistent with a recent evolutionary innovation that may have allowed the *SR* chromosome to escape suppression.

We next used *SweepFinder2* (30) to detect signatures of recent selective sweeps on the *SR* chromosome and found strong evidence of a recent selective sweep in the *Terminal* inversion (Figure 5C). It is important to note that the *Terminal* inversion is estimated to be ∼700,000 years old (26). Our observations of depleted genetic diversity and detection of selective sweeps on *Terminal* likely reflect more recent changes that enable the *SR* chromosome to evade suppression, rather than the initial emergence of the *Terminal* inversion. Taken together, the *D. pseudoobscura SR* chromosome behaves in a manner consistent with an active evolutionary arms race with suppressors, suggesting that changes in the *Terminal* inversion enable the ancestral *BM* form of the driver to evade suppressors. Our results suggest that the *SR* chromosome is shaped by a history of an evolutionary arms race, and that the unsuppressed state the *SR* chromosome represents a transient state awaiting the *de novo* emergence of new suppressors. However, the waiting time for suppression is significant, at least compared to other instances of rapid evolution (31–33).

### Limits to resistance against the *SR* chromosome

The nearly 100% efficient drive mechanism of the *SR* chromosome, found at high frequencies in natural populations, imposes strong selection for the emergence of suppressors. Yet, *D. pseudoobscura* appears to harbor no suppressors of *SR*. We first considered whether this is a sampling problem. With our sampling of over 300 genetic backgrounds, we found zero resistant lines. If we focus on *Y*-linked resistance, this finding leads to a point estimate of 0% Y-linked resistance, with a 95% upper confidence limit of 1.5%. We conclude that it is unlikely that we failed to sample a rare resistant *Y*-chromosome.

Additionally, it is possible that a resistant *Y*-chromosome emerged recently and hasn’t yet swept to a detectable frequency. We estimate that if a resistant *Y*-chromosome were to emerge, it would take less than 100 generations or ∼10-20 years to reach a detectable frequency (See appendix). Because natural populations of *D. pseudoobscura* have been sampled for nearly 90 years, it appears unlikely that a recent resistant *Y*-chromosome is still “taking off”.

There are at least two possible scenarios that may explain the lack of suppression of *SR*. First, under the *gene drive cocktail* scenario, multiple independent drive mechanisms on the *SR* chromosome may target the *Y*-chromosome. We show above that the mechanism of *SR* involves multiple genes in both the *BM* and *T* regions, consistent with the *gene drive cocktail* scenario. In this scenario, resistance would require not a single mutation, but a combination of several resistance mutations to be present simultaneously. This is more problematic for non-recombining *Y* chromosomes than for autosomes. Natural selection would favor *Y*-chromosomes only when they make sons when paired with the *SR* chromosome. Such a *Y*-chromosome must be simultaneously resistant to all drive mechanisms. Under this scenario, the wait time for such a *Y*-chromosome is on the order of (1/*μ*)*^n^*, where *μ* is the mutation rate and *n* is the number of drivers. If we assume *n* = 3 drivers and *μ* = 10^−9^, this yields 10^27^ generations – a long time to wait. Note that this calculation ignores the low probability that any given mutation will become fixed, so the wait time would likely be much longer. Since there is no evidence that the *T* inversion drives on its own, this dampens our enthusiasm for the gene drive cocktail scenario.

Second, the lack of resistance may be explained if the target of drive is essential. Under this scenario, a mutation that removes the target of drive is so costly that it can’t spread even though the *Y*-chromosome evades drive, because it *still* has low fitness. This is unlike the case of *Responder* in *D. melanogaster*, which is targeted by *SD*, but reduction in *Responder* copy number provides immediate and low cost resistance (34). Using established models (5, 6), we found that a *Y*-linked resistance allele should be able to invade and persist in a population even if costly, with costs nearing 30% compared to wildtype *Y*-chromosomes (See appendix). Therefore, the *unditchable target* scenario is possible if *SR* targets something essential for fertility, making resistance alleles extremely costly.

Taken together, these results suggest that the ancient gene drive paradox in *D. pseudoobscura SR* chromosome is explained by a long wait time for the *de novo* emergence of non-costly suppressor alleles or the simultaneous appearance of multiple resistance mutations on a single *Y*-chromosome haplotype. It is important to note that the *gene drive cocktail* and *unditchable target* scenarios are not mutually exclusive, and uncovering the mechanism of drive may distinguish between these possibilities. These findings also suggest that, when resistance arises, its spread would be detectable over human timescales. Repeated sampling of natural populations of *D. pseudoobscura* may position us to catch the emergence of resistance, if it were to arise in the future.

## DISCUSSION

Whether selection is imposed by environmental conditions, biotic interactions, or genetic conflict, the evolutionary response is expected to be rapid. Studies of natural populations have detected many such cases of rapid evolutionary response to selection (31, 35–37). Two examples of rapid evolutionary change imposed by meiotic drivers come from the *SD* system in *D. melanogaster* and the *Paris* system in *D. simulans*. In the *SD* system in *D. melanogaster*, *SD-Mal* – a newer, more efficient driver – recently swept and replaced older, less efficient versions of *SD* (38, 39). In the *Paris* system in *D. simulans*, resistant *Y*-chromosomes are currently sweeping in natural populations in response to the spread of a selfish *X*-chromosome (40). Such systems provide valuable insight into the temporal dynamics of evolutionary arms races with selfish chromosomes.

*D. pseudoobscura SR* has been comprehensively sampled in natural populations over eight decades (10, 12–17). The lack of genetic suppressors against *D. pseudoobscura SR* despite strong selection, sufficient time, and ample genetic variation is surprising. One possible explanation for this observation is that the *tempo* of intra-genomic arms races may slow down with increasing time and complexity of drive systems. When a selfish chromosome first arises, suppressors may be drawn from ample standing genetic variation. Older drive systems may have exhausted all standing natural genetic variation for suppressors, and paths to suppression may thus be more mutationally limited. The old age and complexity of the *D. pseudoobscura SR* system alone, however, does not appear sufficient to explain why natural genetic variation for resistance is absent in this system. For example, *D. melanogaster SD* is also an old and complex drive system, and yet appears to harbor genetic variation for resistant target chromosomes (41). The lack of resistant *Y*-chromosomes in the *D. pseudoobscura SR* system may be better explained if both driver and target of drive are *unditchable*. In *D. simulans*, full expression of the selfish *Distorter-on-X* (*Dox*) gene family is not required for fertility, and suppression occurs through hairpin RNAs that silence these drive genes (42). In *D. melanogaster*, *Responder* repeats (the target of *SD*) can be deleted with little or no observable fitness cost, thus providing immediate resistance (34, 43). In *D. pseudoobscura SR*, if both driver and target of drive are essential, then silencing drive genes or deleting the target of drive do not provide viable paths to restoring Mendelian segregation. The paths to resistance may then be restricted to mutations that are exceptionally rare or costly. A punctuated arms race constrained by mutational limitations may explain the long-term maintenance of this selfish chromosome in an unsuppressed state.

## MATERIALS AND METHODS

### Fly Strains

| Genotype | Purpose | Source |
| --- | --- | --- |
| *SR/se,sh* | Full SR chromosome | Phadnis Lab |
| *BM,sh/se,sh* | BM chromosome | Phadnis Lab |
| *Terminal/se* | Terminal chromosome | Phadnis Lab |
| Y22 | Y-panel | Bachtrog Lab |
| Y10 | Y-panel | Bachtrog Lab |
| Y12 | Y-panel | Bachtrog Lab |
| Y11 | Y-panel | Bachtrog Lab |
| Y25 | Y-panel | Bachtrog Lab |
| Y17 | Y-panel | Bachtrog Lab |
| Y04 | Y-panel | Bachtrog Lab |
| Y15 | Y-panel | Bachtrog Lab |
| Y19 | Y-panel | Bachtrog Lab |
| Y01 | Y-panel | Bachtrog Lab |
| Yref | Y-panel | Bachtrog Lab |
| Y08 | Y-panel | Bachtrog Lab |
| Y21 | Y-panel | Bachtrog Lab |
| Y07 | Y-panel | Bachtrog Lab |
| Y05 | Y-panel | Bachtrog Lab |
| Y02 | Y-panel | Bachtrog Lab |
| Y18 | Y-panel | Bachtrog Lab |
| Y23 | Y-panel | Bachtrog Lab |
| Y03 | Y-panel | Bachtrog Lab |
| Y09 | Y-panel | Bachtrog Lab |
| Y14 | Y-panel | Bachtrog Lab |
| Y20 | Y-panel | Bachtrog Lab |
| Y06 | Y-panel | Bachtrog Lab |
| Ymed | Y-panel | Bachtrog Lab |
| Ylarge | Y-panel | Bachtrog Lab |
| L1 | Wild caught Isogenic Line | Steve Shaeffer |
| L2 | Wild caught Isogenic Line | Steve Shaeffer |
| L3 | Wild caught Isogenic Line | Steve Shaeffer |
| L4 | Wild caught Isogenic Line | Steve Shaeffer |
| L5 | Wild caught Isogenic Line | Steve Shaeffer |
| L6 | Wild caught Isogenic Line | Steve Shaeffer |
| L7 | Wild caught Isogenic Line | Steve Shaeffer |
| L8 | Wild caught Isogenic Line | Steve Shaeffer |

### Collection of wild *D. pseudoobscura* flies

Wild flies were collected near Zion National Park in Southern Utah during September of 2023. 10-20 buckets of bananas were placed at intervals 1-2 miles apart, and wild flies were collected twice per day, once in the morning and once in the evening, when *Drosophila pseudoobscura* are most active. Flies were harvested from buckets of bananas with a Drosophila net (BioQuip; 7110DA). After flies were caught and placed in vials, *Drosophila pseudoobscura* males were separated from other flies, and crossed to D. pseudoobscura *SR*/*se, sh* females. If this cross produced only females or nearly all females, the male was considered an *SR* male and was frozen for sequencing. If the male produced half male offspring, the crossing scheme shown in Figure 1 was conducted to determine whether its *Y*-chromosome was resistant to *SR* drive.

### Crosses to quantify *Sex-Ratio* drive

Males from all *Y*-panels and isogenic strains were crossed to *SR/se,sh* females. These females will produce *se,sh* control males and *SR* test males, both carrying a set of autosomes and *Y*-chromosomes from the *Y*-panel or isogenic strain. From this cross *SR* males appear wildtype, and *ST* males carry the *se,sh* mutations (Figure 1A). All crosses to quantify sex-ratio drive were performed at 23°C. Mating was allowed for 5 days. Offspring were counted until all flies were eclosed.

### Sequencing and genomic analysis

14 *SR* males and 14 *ST* males were frozen after wild collection from Zion National Park, Utah. DNA was extracted from each individual male using Qiagen DNEasy Blood and Tissue Kit. Library preparation and whole genome sequencing and was performed by the University of Utah High-Throughput Genomics (HTG) Shared Resource. Sequencing was done using the Illumina Novaseq X platform. Raw sequencing data can be accessed through the SRA database with accession ID PRJNA1268323. Reads were trimmed using Trimmomatic and aligned using bwa. Analysis of nucleotide diversity was done using ANGSD. PCANGSD was used to perform PCAs. Plots were made using ggplot2. Scripts can be found at https://github.com/JacksonRidges/sr_arms_race. Detection of selective sweeps for *SR XR* was done with SweepFinder2 using the *D. pseudoobscura* MV25 reference genome as the ancestral genome.

## APPENDIX

### Section 1: Invasion time for Y-linked suppressor

It is possible that recently evolved suppressors are slowly increasing in frequency and are still undetectable at the sample sizes employed in the past. To test this, we introduced a suppressor at frequency 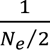 (a single resistant chromosome) and performed deterministic simulations to determine how long it takes for the suppressor to rise from that starting frequency to 5% of all *Y* chromosomes (which we assume would be detected in screens for suppression, including ours). Using the values set out in Larner et al (44), most parameter combinations for the cost of the suppressor and the cost of the driver in males leads to invasion of the *Y*-linked suppressor in less than 100 generations (and often less than 25; Figure A.1).

**Figure A.1.**
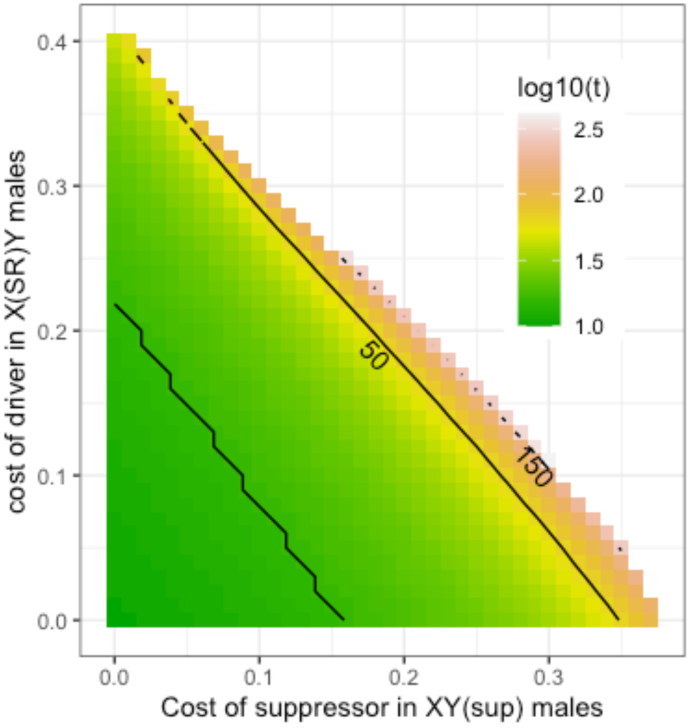
Time (generations) to reach 5%

### Section 2: Invasion of Y-linked suppressor

Next, we explore the parameters that allow for the invasion of a costly *Y*-linked suppressor. As a first pass, we assume a simple population genetic model with perfect *X*-linked drive (all sperm that result in fertile offspring are *X*-bearing), and a rare *Y*-linked suppressor with cost *s_sup_*. In males with both the driving *X* and the suppressing *Y* (*X*^SR^*Y*^sup^), ½+*d_sup_* of the sperm carry the X-chromosome and ½-*d_sup_* are *Y*-bearing. Thus, if *d_sup_* is 0, fair Mendelian segregation of the sex-chromosomes is restored and ½ of the sperm are *X*-bearing. Assume that the driver segregates at frequency *q* in males. The average fitness of a wildtype *Y*-chromosome is:

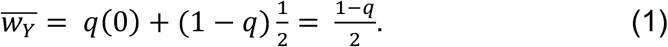

Similarly, the average fitness of a suppressing *Y*-chromosome is:

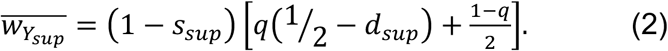

The suppressor can spread when its average fitness is greater than that of the wildtype *Y*-chromosome, which is true when:

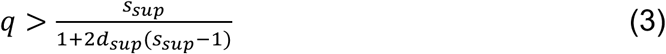

If suppression is perfect (*d_sup_* = 0), this simplifies to *q* > *S_sup_*. Figure A.2 shows this relationship. Note that for a frequency of drive around 0.3 (as for *D. pseudoobscura*), a complete suppressor (as observed in the *BM* population) can be quite costly (*s_SRY_* < 0.3) and still invade. In this case, 70% of the time the suppressor is deleterious because it occurs with a standard *X*-chromosome.

**Figure A.2.**
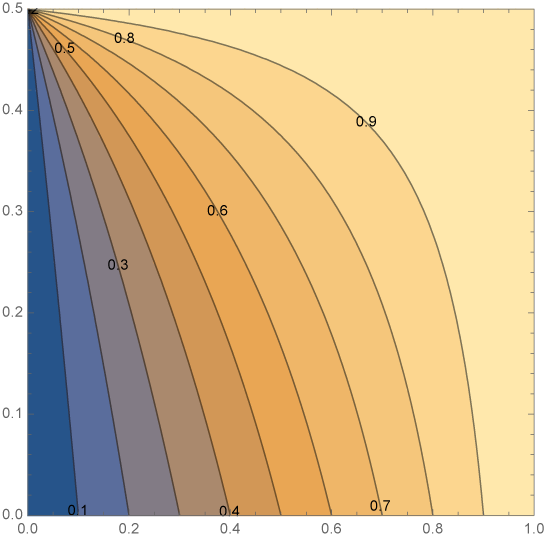
Critical drive frequency for invasion of a suppressing Y with cost *ssup*. X axis is cost of suppressor, Y axis is suppression strength (dsup).

### Section 3: Equilibrium of Y-linked suppressor

Given that the suppressor invades, we next are interested in the equilibrium frequency of such a suppressor. We explore this by building on the model described in Ma *et al.* 2022 (45) which is a modified version of Hall 2004 (5). These modifications allow for a driving *X*-chromosome that results *X^SR^Y* males siring ½+*d* daughters and ½-*d* sons. However, in the presence of a *Y*-linked suppressor, *X^SR^Y^sup^*males sire ½+*d_sup_* daughters and ½-*d_sup_* sons. We restrict both *d* and *d_sup_* to between 0 and ½ and always assume that *d_sup_* is less than *d.* The driving *X* chromosome can carry a host in females (*s_sr_* in *X^SR^X^SR^* females and *h*s_sr_* in *XX^SR^* heterozygous females) or males (*s_sry_*), and the suppressing Y^sup^ carries a cost *s_sup_*.

For our purposes, we will also assume parameters that are in the ballpark of what is expected for *D. pseudoobscura.* Thus, drive is perfect if unsuppressed (*d*=1/2) since all males sired by *SR* fathers are *XO* and therefore sterile. Larner *et al.* (44) estimated the relative of fitness of females heterozygotes for the *SR* chromosome to be 0.92 and homozygotes 0.41 which corresponds to *h* ≈ 0.135 and *s_SR_* ≈ 0.59. They argued that *SR* males have relative fitness of 0.9-1.0 compared to standard males, corresponding to 0 < *s_SRY_* < 0.1. However, polyandry appears to play an important role in limiting the frequency of the driver in *D. pseudoobscura* (16, 18, 46) and for simplicity, we will bake polyandry into the cost of the driver, which allows for a wider range of possible costs (0 < *s_SRY_* < 0.4).

In the absence of a *Y*-linked (or autosomal) suppressor and with perfect drive (*d*=1/2), the equilibrium frequency of the driver is:

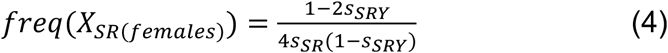

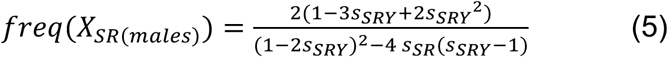

Using the approximations from Larner et al., and a range of costs of the driver in males, the equilibrium frequency of the driver (Eq. 4 and 5) is shown in Figure A.3. Equation 5 suggests that a drive frequency of 0.3 in males corresponds to an effective cost (*s_SRY_*) of about 0.33 in males.

**Figure A.3.**
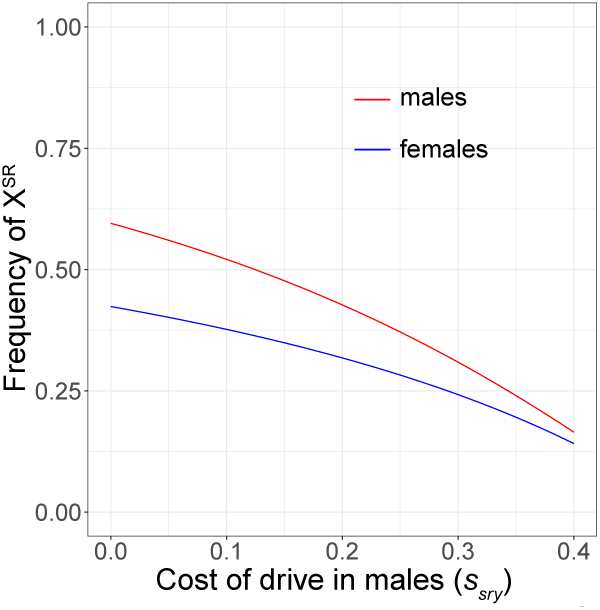
Equilibrium frequency of X^SR^ using parameters from Larner *et al. Blue is frequency in females, red is frequency in males*

We next found the overall solution to the equilibrium given driver and suppressor are segregating. Figure A.4 shows how the equilibrium frequencies of the driver in males and the suppressor are related. This suggests that only a limited parameter space allows for the segregation of the driver at observed frequencies (around 30%) and the suppressor at low (less than 5%) frequency. In fact, for the *Y*-linked suppressor to be found at less than 5% frequency and the driver to be between 10 and 40% in males, there must be high cost of drive in males and low cost of suppression or lower cost of the driver, but intermediate cost of the suppressor (Figure A.5).

**Figure A.4.**
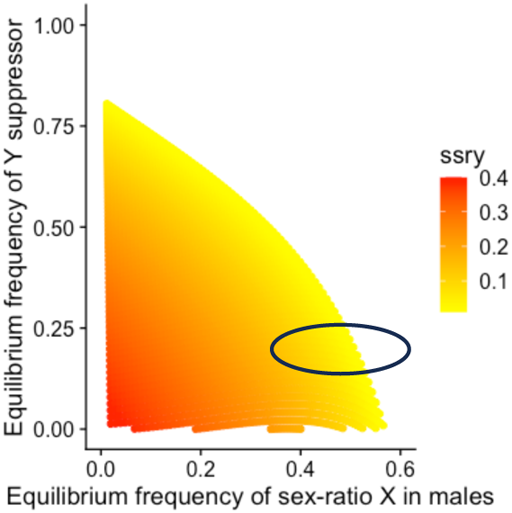
Equilibrium frequency of X^SR^ and Y^sup^ using parameters from Larner *et al.* We expect that, if suppressors segregate at equilibrium in *D. pseudoobscura*, that parameter space is in the area circled in blue.

**Figure A.5.**
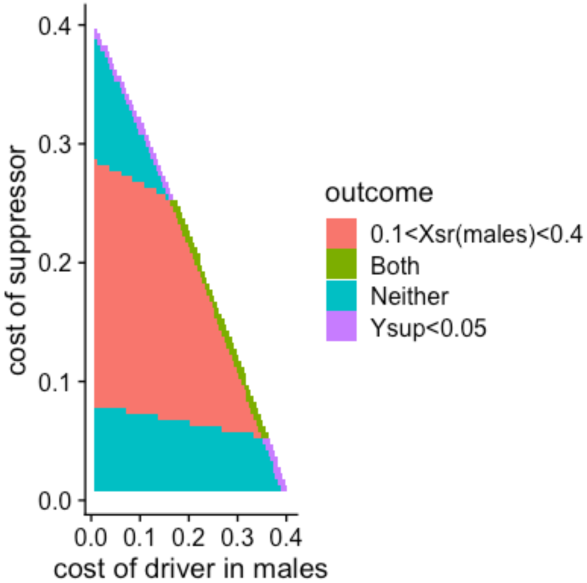
Conditions for low frequency suppression and intermediate frequency of drive in males. Note, white space above the diagonal leads to the loss of the suppressor.

## ACKNOWLEDGEMENTS

We thank Matt Hahn for helpful discussions. We thank Steve Schaeffer and Doris Bachtrog for providing fly strains. This work was supported by the National Institute of Health grants R35GM156267 to NP, 5T32GM141848 to JB, and NSF MCB CAREER grant 2047052 to RLU.

---

**Supplemental Figure 1:**
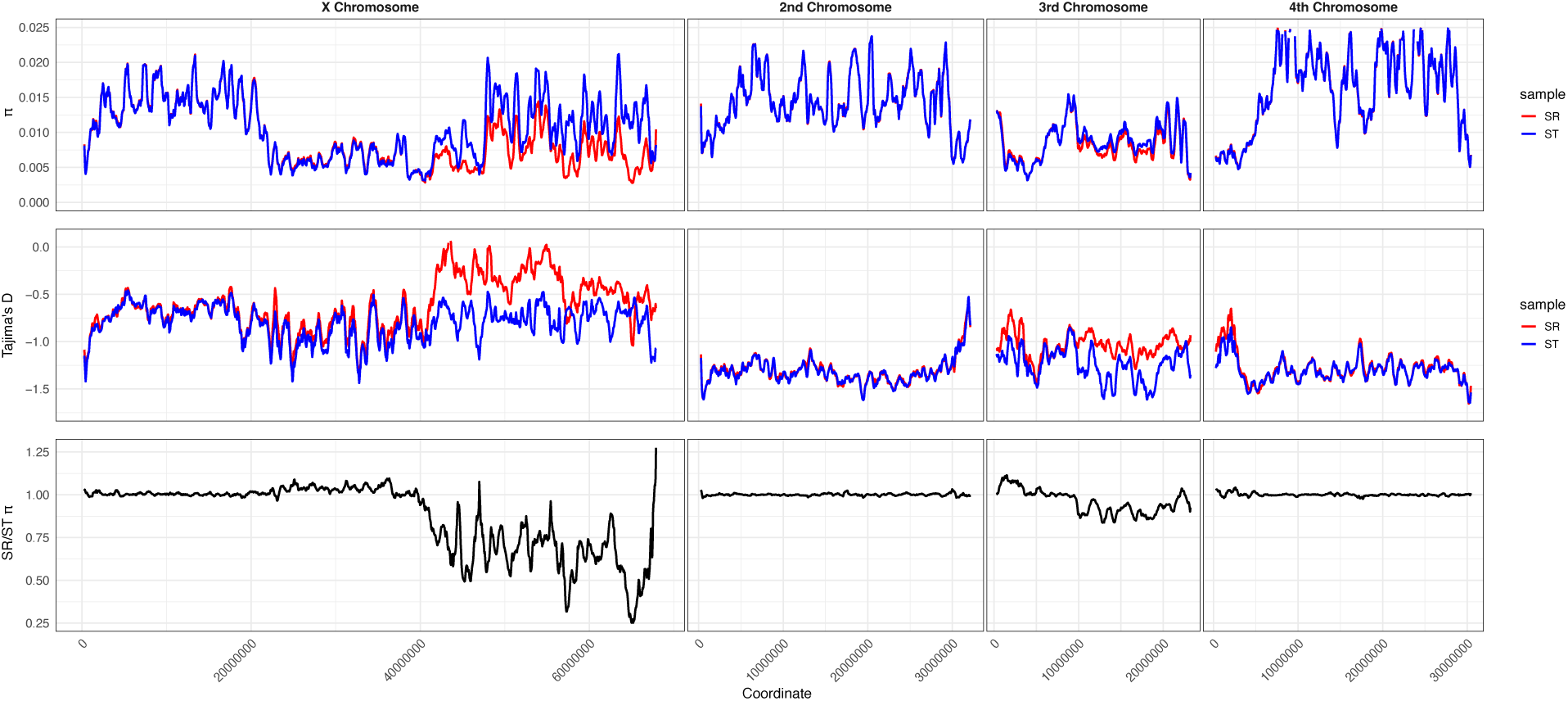
π, Tajima’s D, and *SR*/*ST* π ratio across all chromosomes for 28 individually sequenced *ST* and *SR* males.

**Supplemental Figure 2:**
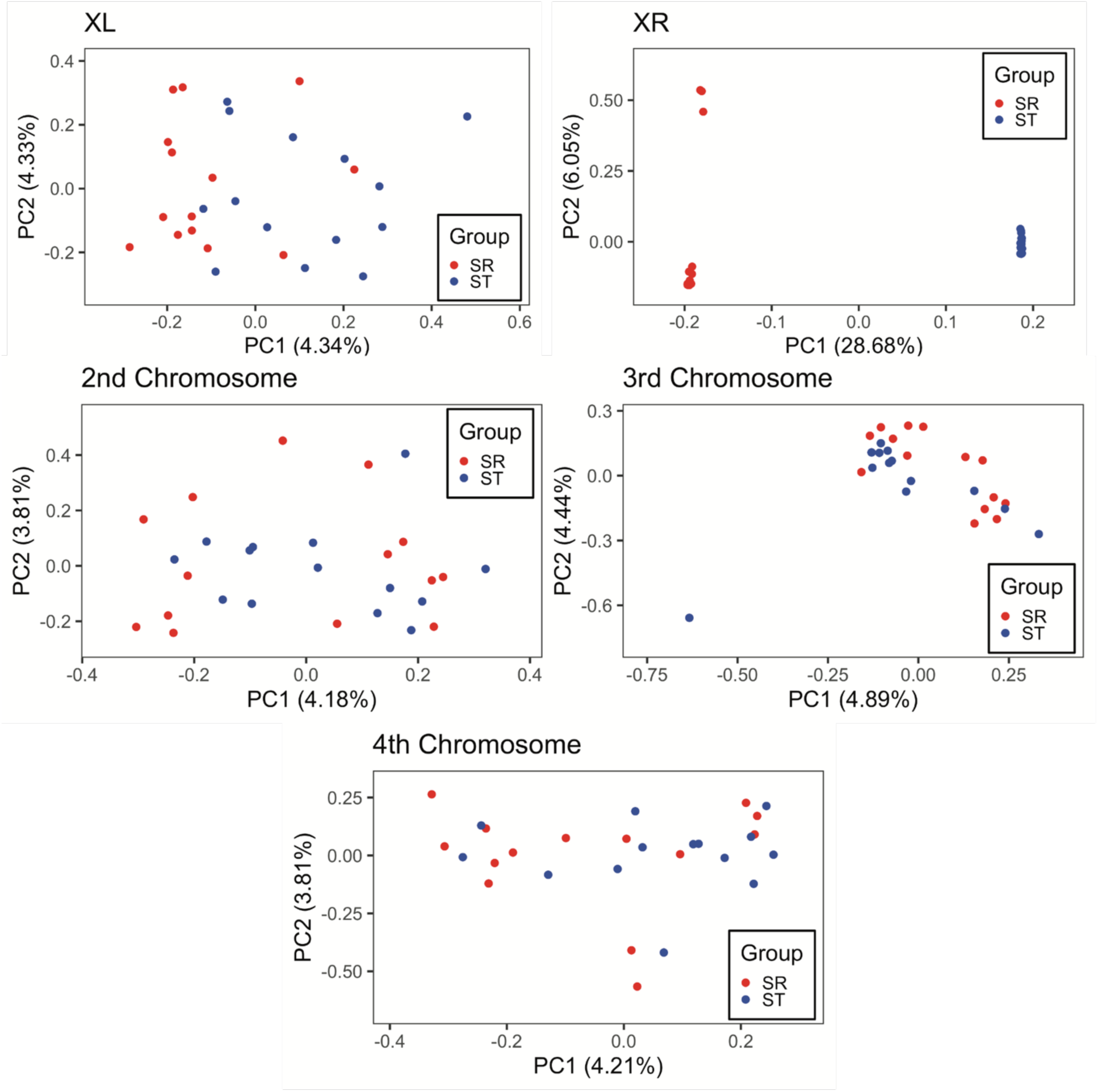
Principal Component Analysis for all chromosomes generated by PCANGSD.

